# Immunohistochemical assays for bladder cancer molecular subtyping: Optimizing parsimony and performance using Lund taxonomy

**DOI:** 10.1101/2021.08.06.455345

**Authors:** Céline Hardy, Hamid Ghaedi, Ava Slotman, Gottfrid Sjödahl, Robert J. Gooding, David M. Berman, Chelsea L. Jackson

## Abstract

Transcriptomic and proteomic profiling reliably classifies bladder cancers into luminal and basal molecular subtypes. Based on their prognostic and predictive associations, these subtypes may improve clinical management of bladder cancers. However, the complexity of published subtyping algorithms has limited their translation into practice. Here we optimize and validate compact subtyping algorithms based on the Lund taxonomy. We reanalyzed immunohistochemistry (IHC) expression data of muscle-invasive bladder cancer samples from Lund 2017 (n=193) and 2012 (n=76) cohorts. We characterized and quantified IHC expression patterns, and determined the simplest, most accurate decision tree models to identify subtypes. We tested the utility of a previously published algorithm using routine antibody assays commonly available in surgical pathology laboratories (GATA3, KRT5 and p16) to identify basal/luminal subtypes and to distinguish between luminal subtypes, Urothelial-Like (Uro) and Genomically Unstable (GU). We determined the dominant decision tree classifiers using four-fold cross-validation with separate uniformly distributed train (75%) and validation (25%) sets. Using the three-antibody algorithm resulted in 86-95% accuracy across training and validation sets for identifying basal/luminal subtypes, and 67-86% accuracy for basal/Uro/GU subtypes. Although antibody assays for KRT14 and RB1 are not routinely used in pathology practice, these features achieved the simplest and most accurate models to identify basal/luminal and Uro/GU/basal subtypes, achieving 93-96% and 85-86% accuracies, respectively. When translated to a more complex model using eight antibody assays, accuracy was comparable to simplified models, with 86% (train) and 82% (validation). We conclude that a simple immunohistochemical classifier can accurately identify luminal (Uro, GU) and basal subtypes and pave the way for clinical implementation.

## Introduction

Molecular profiling of bladder cancers has provided insight into the biology and clinical behaviour of tumors. In muscle-invasive bladder cancer (MIBC), work pioneered by several groups characterized the genomic and proteomic profiles of these tumors to identify molecular subtypes (1–4). This work has yielded a series of refined and overlapping subtyping taxonomies. At the top hierarchical level and common to all of these subtyping schemes is the identification of luminal-like and non-luminal-like (basal-like) subtypes, representing divergent tracks of urothelial differentiation (2,5). Thus, the various subtyping taxonomies were independently developed, but now converge in a consensus subtyping scheme that identifies basal and luminal subtypes, and can be variably subclassified into 3-7 groups (1–6). However, this model has yet to be validated at the protein expression level.

The Lund bladder cancer research group was among the first to establish genomic subtyping, initially relying on mRNA (1). Noticing the shortcomings of transcriptomic profiling, they validated mRNA-based subtyping at the protein expression level, specifically in cancer cells, using immunohistochemistry (IHC) (7, 14–16). This work clarified five main tumor cell-intrinsic subtypes (tumor cell phenotypes) of MIBC: Genomically Unstable (GU), Urothelial-like (Uro), Basal/Squamous Cell Carcinoma-like (Basal/SCCL), Mesenchymal-like (Mes-like) and small-cell/neuroendocrine-like (NE-like). These subtypes reflect the biological heterogeneity of MIBC tumors through a variety of pathogenic mechanisms. For example, luminal subtypes (Uro, GU) express markers of luminal differentiation programming, such as GATA3, FOXA1, and PPARy (26–28). Conversely, the basal subtype expresses basal keratins, such as KRT5 and KRT14 (26–28). Notably, the distinction between the two luminal subtypes, Uro and GU is not universal to all subtyping schemes but is marked by inverse protein expression patterns involving key alterations in RB1, FGFR3, CCND1 and CDKN2A.

Importantly, these subtypes have demonstrated differences in clinical outcomes where broadly, tumors of the basal subtypes are aggressive and have poor prognosis, while luminal subtypes are associated with increased overall survival (1–7). Work from the Lund group has identified further prognostic differences amongst the luminal subtypes, such that the GU subtype experiences a worse prognosis relative to its Uro counterpart. Molecular subtypes have also been associated with response to neoadjuvant chemotherapy and immune checkpoint blockade (3, 8–13). These predictive and prognostic associations indicate potential clinical applications of subtyping. However, subtyping has yet to be translated to the clinical setting. The complexity of transcriptomic profiling methods and variable sampling of tumor and stromal (benign supportive or immune cells) have hindered the impact of this work (14). These challenges have prevented the development of a single, consistent, and simple methodology to elucidate the clinical implications of molecular subtyping.

Given its simplicity and availability in routine clinical pathology labs, IHC may serve as an attractive, simple alternative to transcriptomic profiling. Indeed, recent work has focused on using IHC to identify luminal and basal subtypes (10, 18–20). Multiple studies identified key immunohistochemical protein features capable of achieving high accuracy for identifying mRNA-subtypes (10, 18–20). However, to our knowledge this work has yet to extend beyond luminal/basal subtyping and has focused on validating mRNA subtypes as opposed to the tumor cell phenotypes (IHC subtypes) clarified by the Lund taxonomy. The Lund group tested a comprehensive panel of 24 key IHC features that can accurately classify bladder cancers into the five Lund subtypes (7,14). Based on antibody availability and staining reproducibility, they further distilled these to a set of 13 core IHC assays (7,14). In contrast to other subtyping schemes, the Lund taxonomy has the highest resolution classification scheme, bears the closest similarity to the consensus model, and has been the most extensively validated using IHC.

We recently proposed a more concise model using antibodies against GATA3, KRT5 and p16 to identify the three main subtypes of MIBC, which account for over 90% of all samples (21; Jackson 2021). Furthermore, it utilized three readily available IHC assays widely used in surgical pathology laboratories, suggesting a simple and accessible algorithm for use across clinical centres (22–25). Since then, new data enabled us to test the accuracy of this algorithm and compare it to an array of algorithms derived from all 13 core IHC assays (14–16). Using these data, we characterized expression patterns and the extent of overlap of core IHC markers between key Lund subtypes (basal/Uro/GU). We then systematically analyzed potential combinations of IHC assays to create simple and accurate subtyping classifiers and compared their performance to the GATA3/KRT5/p16 model.

## Materials and Methods

### IHC staining, score calculations and subtype definitions

Two datasets representing work previously published in 2012 (Lund 2012 dataset) and 2017 (Lund 2017) were assessed (1,14). Images were obtained from sections previously stained (14) with antibodies against key subtyping proteins CCNB1, CCND1, CDH1, CDH3, CDKN2A (p16), CHGA, E2F3, EPCAM, FGFR3, FOXA1, GATA3, KRT5, KRT14, KRT20, NCAM1, PPARG, RB1, RXRA, SYP, TP63, TUBB2B, UPK3, VIM or ZEB2. IHC subtype assignment was determined by calculating a definition (genomic circuit) score, using 13 key immunohistochemical features to assign samples to Uro, Basal, GU, NE-like and Mes-like subtypes, as previously described by Sjödahl et al (2017) (14). IHC phenotype and circuit score definitions are described in Supplementary Methods (Table S1). These IHC subtypes were used as class labels in all subsequent analyses.

### Data scaling and exclusions

Data was available for a total of 327 MIBC samples (1,14). Of these, 193 samples had complete expression data for all 24 proteins. Due to the small number of Mes-like and SC/NE-like tumors (n=16), model building was restricted to identify luminal and basal subtypes, where an adequate number of samples were available to train and validate classifiers. A total of 177 samples of basal, Uro and GU subtypes from the 2017 Lund dataset (14) were included and used for further analysis. Descriptions of evaluation and original scaling of these proteins are described in Supplemental Material (Table S1). As previously described (16), samples were assigned one of five KRT5 expression patterns or proximity scores (all, diffuse, multiple layers, 1-layer, or no staining. KRT5 score values were calculated as the proximity score multiplied by the percentage of positively stained tumor cells (16), as summarized in Table S1. All expression values were scaled to yield a tumor cell score (TCS) from 0-1 in order to interpret IHC expression values on the same scale.

### Data visualization

Statistical tests were performed in R version 3.5.3. Supervised hierarchical clustering and Spearman correlation were performed to visualize expression patterns of 24 proteins across all 193 samples. In order to investigate the shape of the data and identify groups of similar samples, we performed principal component analysis (PCA) and visualized the results using the “FactoMineR” package. In PCA analysis, expression of the 24 available proteins was used as features to identify basal, Uro, and GU samples (n=177). NE-like and Mes-like samples were excluded from PCA analyses due to small sample size (n=16). To select proteins (features) to make a subtype classifier, a series of binary receiver operating characteristic curve analysis (ROC) were performed for each of the 24 proteins using the “pROC” package in the R environment.

### Model Building

We assessed the utility of GATA3, KRT5 and p16, as well as the simplest and most accurate combination of antibodies from all IHC features. The Rpart package was used to perform model building and trees were pruned to the minimum number of nodes to prevent overfitting. Robustness of models was assessed by comparing the performance of trained models to their accuracy on a separate validation set. Four-fold cross-validation with uniformly distributed train (75%) and validation (25%) sets was used to determine the dominant tree structure, which was defined as the model structure which emerged most frequently from cross validation. Accuracy was reported rather than balanced accuracy due to low sample numbers in the basal subtype which overweighs the classification inaccuracy.

To build and visualize a multiclass decision tree incorporating a larger number of proteins, but with optimized accuracy, we used “sklearn” and “graphviz” libraries in python. The dataset was split into train (75%) and validation (25%) datasets. To train the model and at the same time to avoid overfitting, we followed a stratified K-fold cross validation (K = 10) approach. Balanced accuracy of the mean and its standard error of mean (SE) were used to evaluate model performance. Optimal tree depth was determined by running a K-fold cross validation (K = 5). The “entropy” score was considered as a measure of impurity in binary classification.

## Results

### Visualizing protein expression, quantifying redundancy and identifying subtype-defining features

#### Characterizing expression patterns

Using supervised hierarchical clustering of all MIBC samples (n=193) we visualized the expression of 24 IHC features in each subtype. We observed the expected associations of subtypes with key molecular features (Fig. 1A). Luminal (Uro and GU) tumors were characterized by distinctly high expression of GATA3, EPCAM, and FOXA1. Basal tumors showed enhanced expression of basal keratins, KRT5 and KRT14. Spearman correlation of IHC expression values quantified the extent of the overlap of these features, where proteins characteristic of luminal subtypes such as GATA3 and FOXA1 showed strong positive correlation (r = 0.6) as did features of basal differentiation, KRT5 and KRT14 (r = 0.69) (Fig. 1B). Protein expression also reflected underlying subtype-specific pathogenic mechanisms, where RB1 and CCND1 expression levels were positively correlated (r = 0.59) and RB1 and p16 were negatively correlated (r = −0.5).

**Figure 1.**
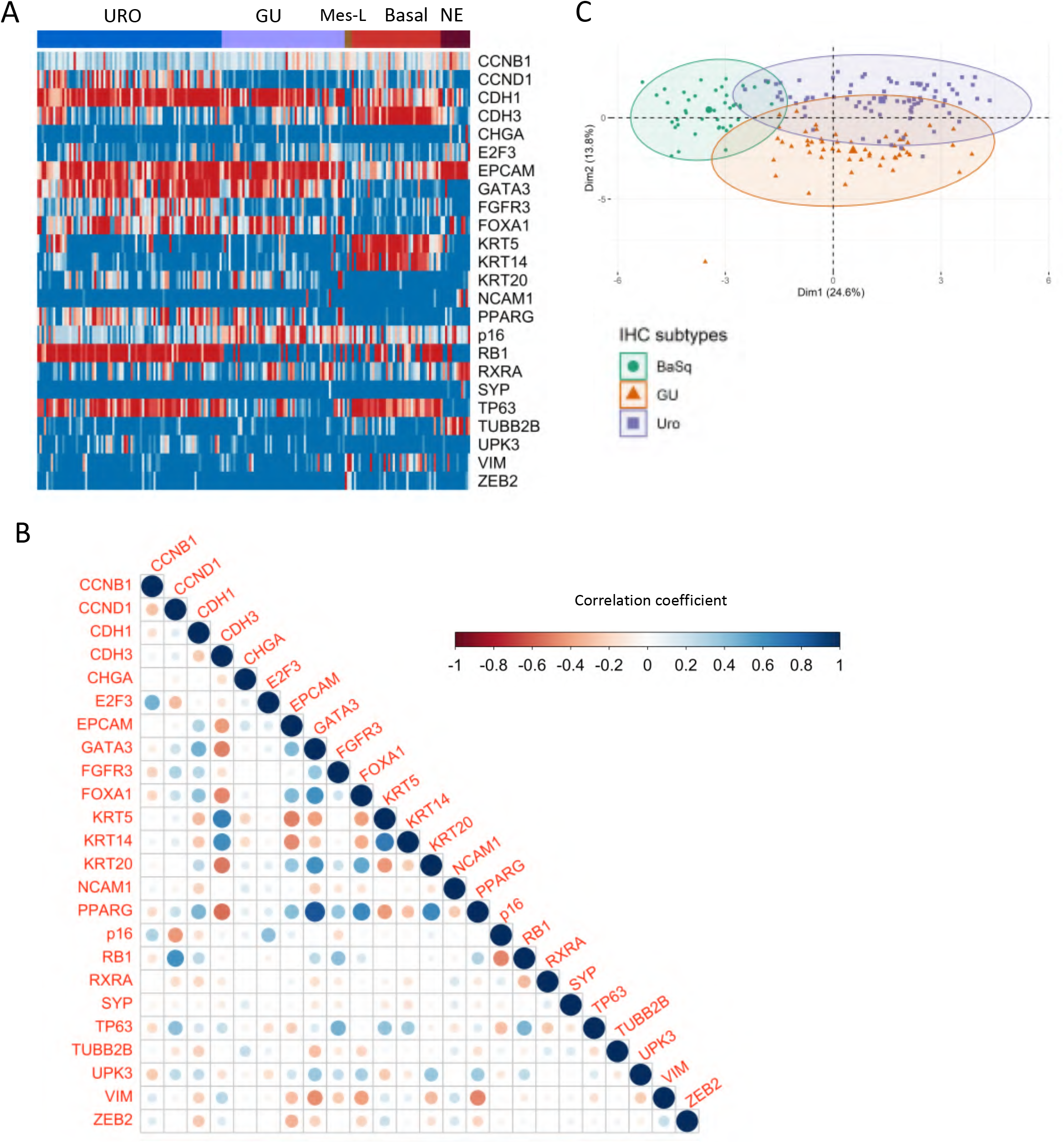
Redundancy of protein features and their relationships with basal and luminal subtypes. (A) Supervised clustering of IHC expression values recapitulates expected patterns defining tumor cell phenotypes of IHC subtypes (B) Spearman correlation of IHC expression for 24 proteins in all samples (n=193). Dot size corresponds to correlation strength. (C) PCA analysis incorporating IHC expression data for Uro, GU and basal subtypes.

Using PCA, we found that similar samples (members of a specific subtype), clustered together. However, there was also overlap between subtypes, which was most extensive between the two luminal subtypes (Uro and GU) (Figure 1C). Interestingly, as previously demonstrated in mRNA profiling as “urobasal b” phenotype (1), the basal cluster overlapped more with Uro than with the GU subtype. These observations highlight redundant functions of several proteins in molecular subtyping and suggest an opportunity to build more compact classifiers with fewer features.

#### Clarifying subtype-defining features

Using binary ROC analysis, we ranked the association of individual proteins with luminal/basal classes and with individual Lund subtypes (basal/Uro/GU). We identified KRT14, KRT5, CDH3, FOXA1, GATA3, PPARG, RB1, CCND1, CDKN2A (p16), FGFR3, and TP63 as the most effective features for classifying subtypes with greater than 80% accuracy in binary ROC analyses (Table 1). As expected, the same key proteins which were previously selected for defining tumor cell phenotypes using the Lund 2017 data (14) achieved the highest accuracies. For discriminating between basal and luminal subtypes, basal keratins ranked highest, with accuracies of 94.39% (95% CI: 91-97%) for KRT5, and 94.11% (95% CI: 89-99%) for KRT14. The best proteins for discriminating between Uro and GU subtypes were RB1 (92.62%, 95% CI: 87-98%), CCND1 (92.62%, 95% CI: 87-98%), and p16 (92.62%, 95% CI: 87-98%) (Table 1).

**Table 1.**
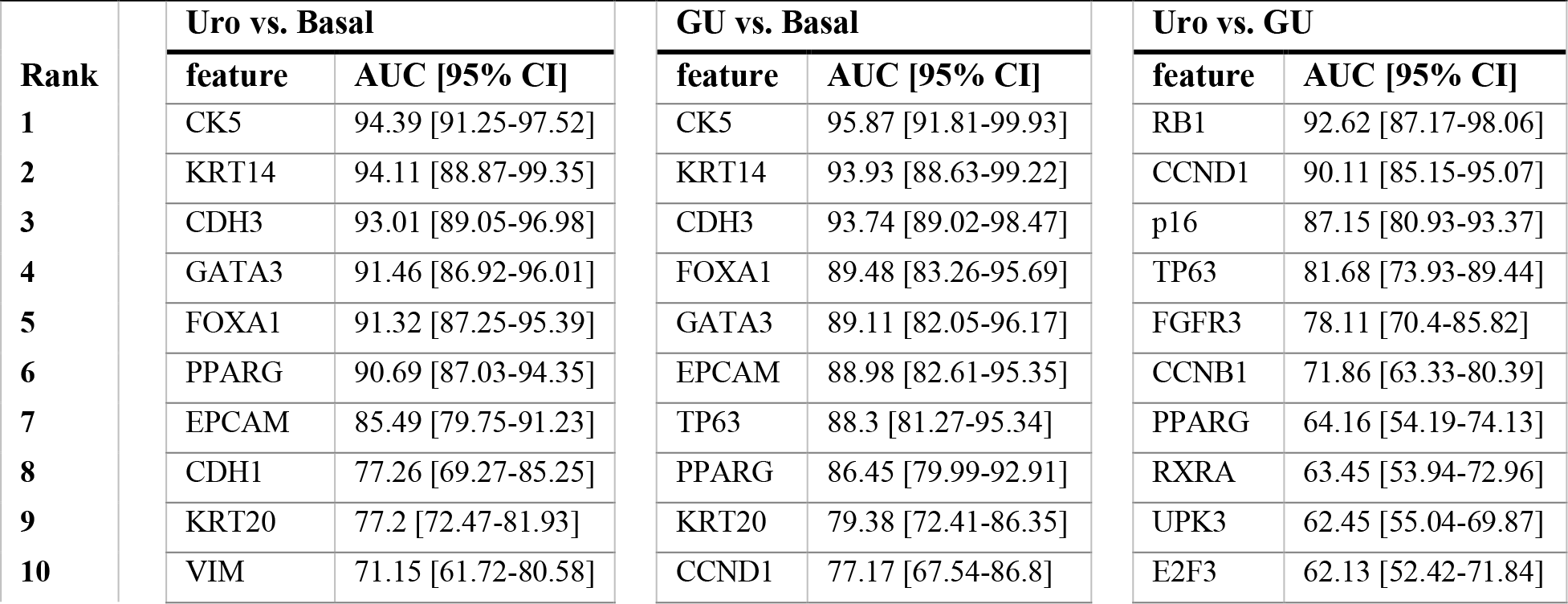
Ranked performance of the top 10 proteins for identifying subtypes in ROC analysis.

Importantly, several proteins were highly correlated with each other (Fig. 1B) and with a given subtype, and could individually identify either basal (KRT14, KRT5) or luminal (CDH3, FOXA1, and GATA3) tumors with high accuracy. These observations suggested that an accurate subtype classification model can be built with fewer protein features. Complete data of ranked accuracies of all 24 proteins is available in Supplementary Table S2.

### Determining the most parsimonious models to accurately identify subtypes

#### Basal/Luminal classification using GATA3 and KRT5

Based on a model we proposed previously (21), we investigated the ability of GATA3, KRT5 and p16 to identify basal and luminal subtypes. Others have reported that the combination of GATA3 and KRT5 can identify basal and luminal subtypes with over 90% accuracy (17,18). Using available Lund datasets of samples subtyped by both mRNA expression and IHC (1,14), we explored the utility of GATA3 and KRT5 for top-level basal/luminal classification. Four-fold cross-validation of a decision tree classifier using uniformly distributed training sets (n=133) identified a dominant tree with three branches that used GATA3 followed by KRT5 (Fig. 2). These branches first separated tumors positive for GATA3 into the luminal branch, following which high expression of KRT5 distinguished basal tumors, and tumors with low expression of both features were classified as luminal (Fig. 2A) Across trees with the dominant structure, a threshold of 0.21 was consistently chosen for GATA3, whereas KRT5 demonstrated a range of thresholds from 0.09-0.34, with high resulting accuracies in separate validation sets. We then sampled iteratively across 100 random validation sets to determine the KRT5 threshold with the highest resulting accuracy. The tested thresholds included those provided by the resulting dominant trees (e.g., 0.09, 0.16, 0.34) as well as any intermediate values within 0.05 increments. Using these 100 validation sets, we calculated the mean accuracy for each threshold and visualized the distribution of accuracies (Table 2). For the range of KRT5 thresholds between 0.09-0.34, luminal/basal classification rates remained highly accurate at 93-94%.

**Table 2.**
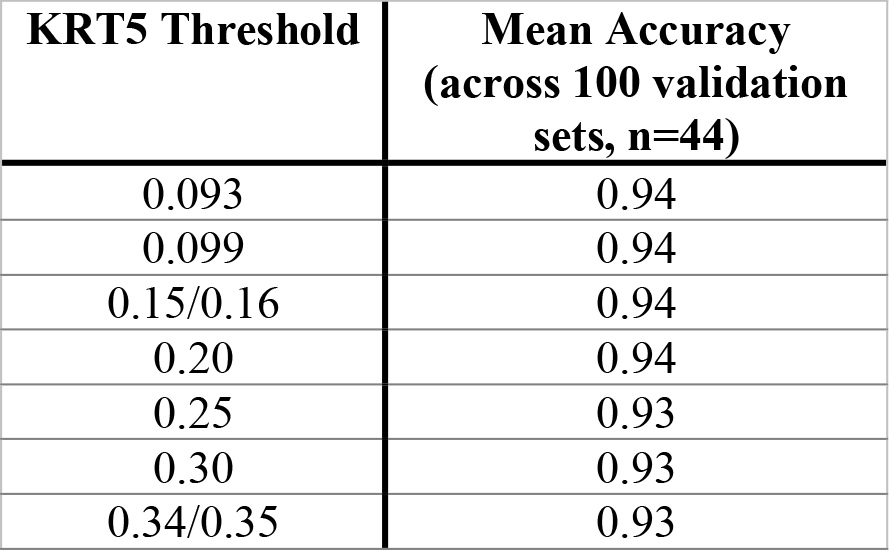
Mean accuracies for identifying Basal and luminal subtypes across 100 random validation sets, using varying KRT5 thresholds.

| KRT5 Threshold | Mean Accuracy<br>(across 100 validation<br>sets, n=44) |
| --- | --- |
| 0.093 | 0.94 |
| 0.099 | 0.94 |
| 0.15/0.16 | 0.94 |
| 0.20 | 0.94 |
| 0.25 | 0.93 |
| 0.30 | 0.93 |
| 0.34/0.35 | 0.93 |

**Figure 2.**
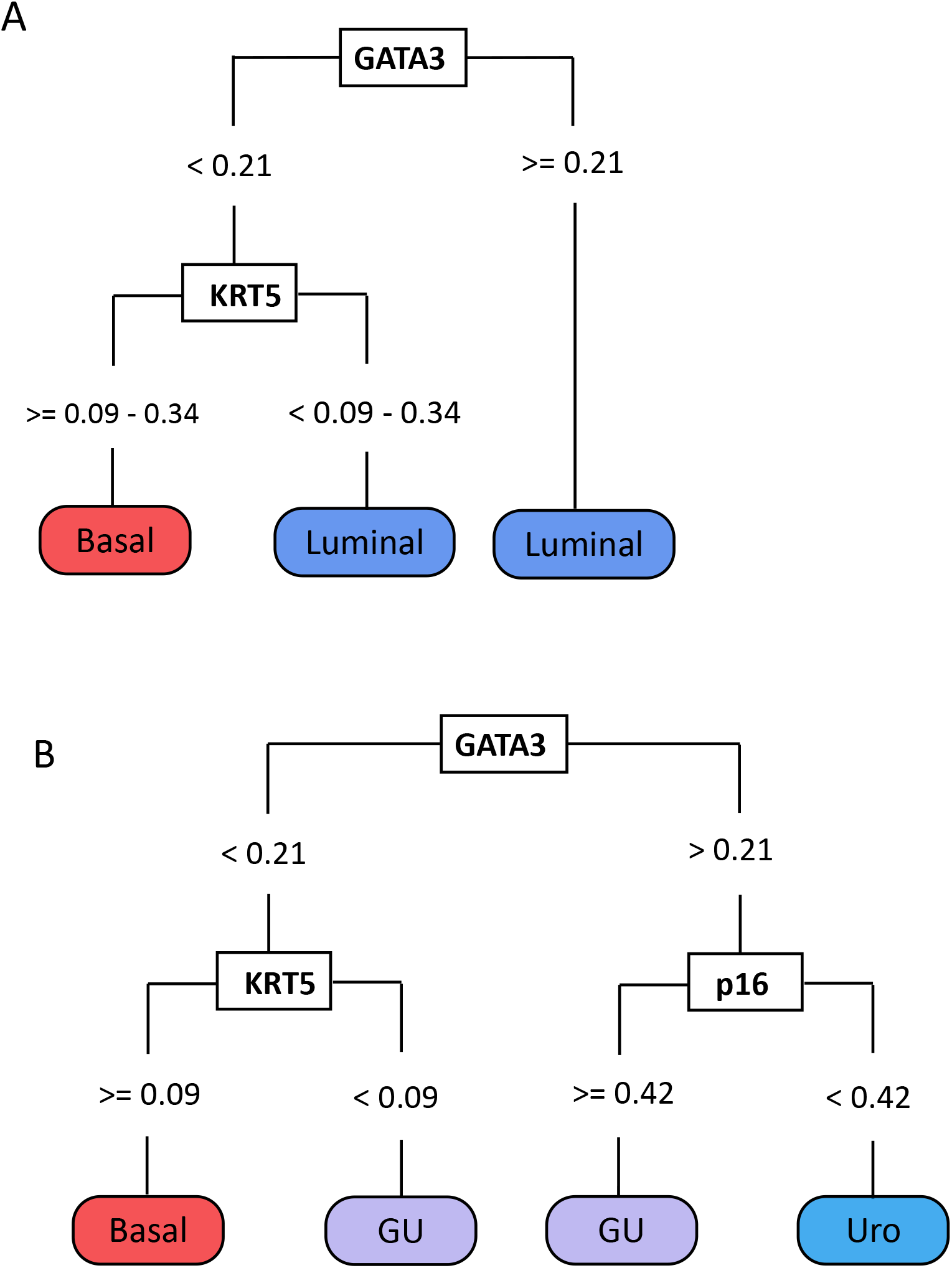
Dominant decision tree classifiers use GATA3, KRT5, and p16 to identify luminal-basal and Uro, GU and basal subtypes. (A) Identification of luminal and basal subtypes using GATA3 and KRT5. (B) Identification of basal, Uro and GU subtypes using GATA3, KRT5 and p16.

To further test the classification accuracy of this model, we used data from two separate cohorts (Lund 2017 (14) and Lund 2012 (1)) as training and validation sets, respectively. This resulted in the same tree structure of GATA3, followed by KRT5, with similar thresholds (0.21 and 0.099 respectively), and an accuracy of 91% (95% CI: 82-96%) (Fig. 2). As a result, we conclude that GATA3 and KRT5 can classify MIBC samples into Luminal and Basal subtypes with >90% accuracy.

#### Uro, GU and basal subtype classification using GATA3, KRT5 and p16

Next, we assessed the utility of a model incorporating p16 to identify Uro and GU subsets of luminal tumors. Similar to our methods for basal/luminal classification, we used the 2017 Lund cohort (14) (n=177) as a training set and an additional MIBC Lund cohort (1) (2012 cohort, n=76) as a validation set. The resulting tree demonstrated four branches. The first node separated tumors positive for GATA3 into the luminal branch, following which high expression of p16 distinguished GU from Uro tumors. This tree suggests that tumors with low expression of both GATA3 and KRT5 are classified as GU, while tumors negative for GATA3, but with expression of KRT5 are basal. The accuracy of the dominant model using GATA3, KRT5, and p16 was 78% (95% CI: 67%-86%) in the validation set. Importantly, using this approach revealed similar thresholds for GATA3 (0.21) and KRT5 (0.099) used in basal/luminal classification and initial cross validation, indicating that these thresholds are likely robust for classification. Therefore, we conclude that using all three antibodies can classify MIBC samples into basal/Uro/GU subtypes with ~ 78% accuracy.

### Optimizing decision-tree models for parsimony and accuracy

#### Basal/Luminal classification using KRT14

We next expanded our analysis to all 24 IHC features to determine which protein features could improve or achieve better classification accuracy. Using the same four-fold cross-validation methods on uniform train and validation sets, we identified a dominant tree structure that separated two branches using only KRT14 (threshold= 0.69) (Fig. 3). This tree classified tumors with high expression of KRT14 (TCS > 0.69) as basal, and tumors with low expression of KRT14 (TCS < 0.69) as luminal. Interestingly, this KRT14 threshold (0.69) was identical across all training cross-validation iterations and therefore did not require additional bootstrapping to optimize thresholds with the highest accuracy. Throughout cross-validation iterations, the accuracy of this model was very high, ranging from 93-96% (95% CI: 0.79-0.99). We were unable to test the accuracy of this dominant tree structure in independent validation sets (2017 cohort) due to differing KRT14 assessment methods between the 2012 and 2017 cohorts (1,14). However, these findings suggest that KRT14 can provide more accurate identification of basal and luminal subtypes than KRT5, but additional testing is warranted to identify its overall accuracy across multiple cohorts.

**Figure 3.**
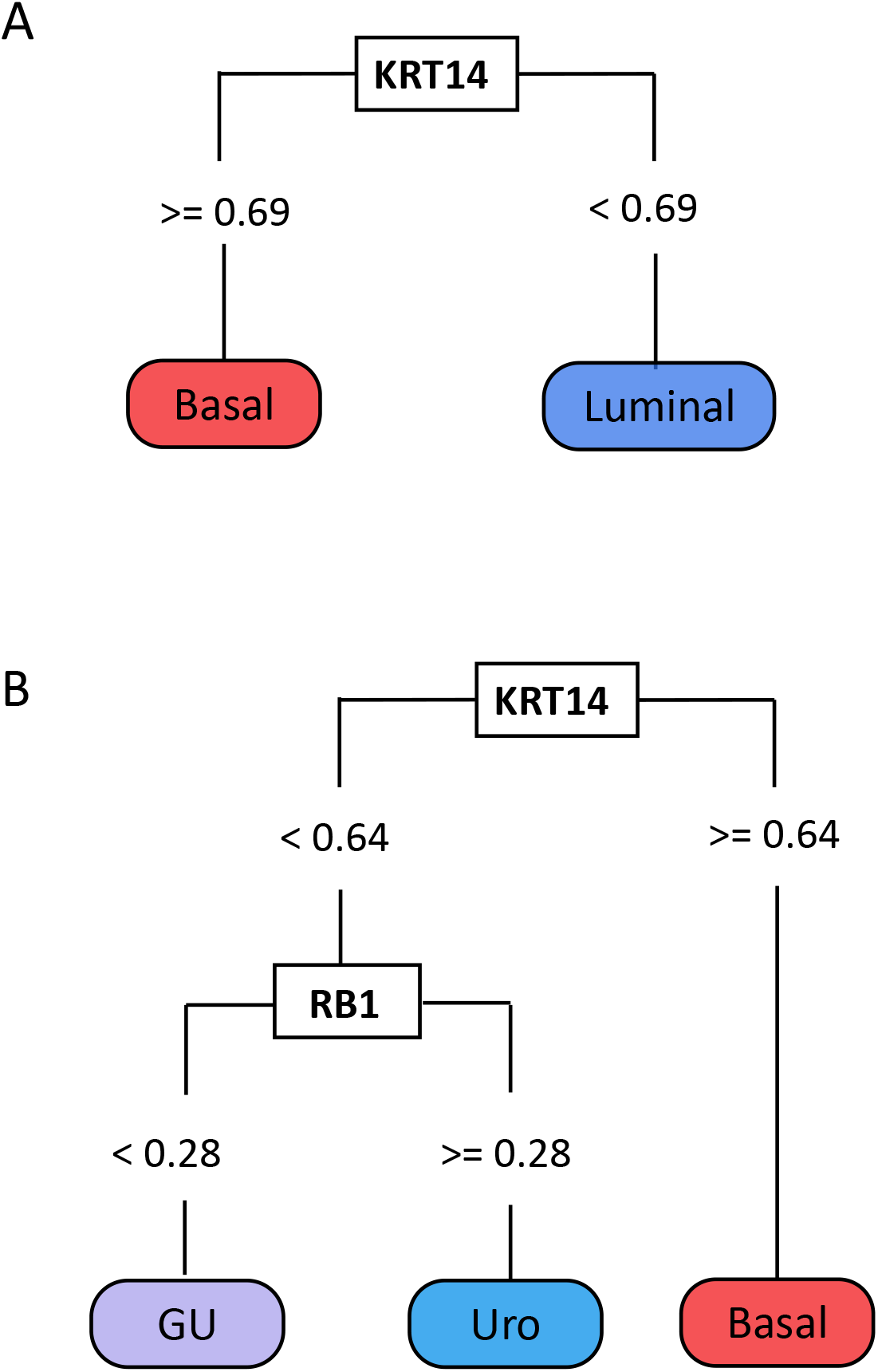
Dominant decision tree classifiers for identifying luminal-basal and Uro, GU and basal subtypes. (A) Identification of luminal and basal subtypes using KRT14. (B) Identification of Uro, GU and Basal subtypes using KRT14 and RB1.

#### KRT14 and RB1 for classifying Basal, Uro and GU subtypes

In building models that classify tumors into the three dominant MIBC subtypes (Basal, Uro, and GU), we identified several models that used only two proteins, KRT14 (threshold = 0.64) followed by RB1 (threshold = 0.28), and achieved an accuracy of 85-86% (95% CI: 0.71-0.94) (Fig. 3). This model classifies tumors that show high KRT14 expression as basal, and low expression as luminal. Luminal tumors are then subdivided into Uro and GU subtypes based on positive and negative expression of RB1, respectively. KRT14 was assessed differently in the 2017 and 2012 cohorts (1,14), thus preventing further examination of these models in independent validation sets. However, the promising accuracies of simplified models using KRT14 and RB1 suggest that these proteins may serve as useful subtyping features.

### Improving accuracy with additional protein features

To assess the performance of more complex classifiers, we made multi-class decision tree classifiers optimized to identify basal, Uro and GU tumors. For feature selection, we considered proteins with area under the curve (AUC) ≥ 80% in ROC analysis for classifying samples into different subtypes. Using these more expansive parameters, the most accurate decision tree incorporated KRT5, KRT14, GATA3, FOXA1, EPCAM, TP63, RB1 and CCND1 into four nodes (Fig. 3). Following 10-fold stratified cross-validation, this model achieved a balanced accuracy of 85.6 % ±0.07 in the training set. When applied to the validation set, the accuracy decreased slightly to 82%. The performance metrics of the model are summarized in Table 4. Interestingly, this model suggested a top-level division based on RB1 expression which primarily serves to separate Uro and GU subtypes. It further selected CCND1 to distinguish GU tumors. However, CCND1 expression in this model served only to classify a small number of samples (one GU from basal, four GU from Uro). Similarly, incorporation of GATA3 into the model distinguished one GU sample from Uro. TP63 was incorporated into the model since GU tumors showed low expression compared to Uro. These observations indicate that the majority of Uro and GU samples are effectively split by the top-level division using RB1, and that the third and fourth level branches provided little value.

**Table 3.** Summary table of decision tree classifier accuracies

|  | Basal/Luminal | Uro/GU/Basal |
| --- | --- | --- |
| **Best simplified model** | **KRT14**<br>93-96 %<br>(79-99%) | **KRT14 +RB1**<br>85-86%<br>(71-94%) |
| **Clinically available assays** | **GATA3 + KRT5**<br>91%<br>(82-96%) | **GATA3 + KRT5 + p16**<br>78%<br>(67-86%) |

**Table 4.**
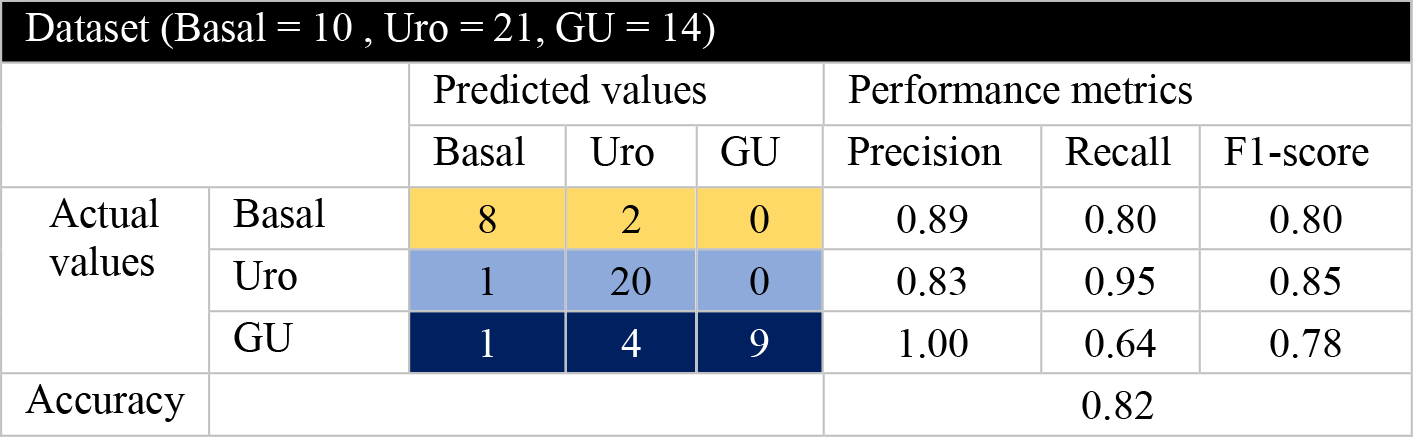
Performance statistics of the optimal classifier model on the validation dataset (n = 45)

### Visualizing IHC staining and distributions of protein expression

#### Distributions of IHC expression

Ideally, biomarkers selected for clinical use would show clear and subtype-specific differences in expression levels, and pathologists would be capable of recognizing these differences without relying on computerized quantitative analysis. To address these concerns, we visualized the distributions of IHC expression across samples. For GATA3, KRT14, KRT5, and RB1, our analysis revealed binary expression patterns, indicating a rational threshold for characterizing positive and negative expression (Fig. 5). This distribution was markedly different in luminal and basal subtypes for KRT14 and KRT5, while GATA3 expression was slightly more variable but still indicated clear separation. On the other hand, p16 expression showed greater variability with a large number of samples showing intermediate staining. While overall, p16 staining appears normally distributed, this is primarily between basal and Uro, or basal and GU subtypes. Since p16 is proposed as a protein feature for the separation of Uro and GU subtypes, where the distribution appears bimodal, this feature is likely still useful for their separation. CCND1 also showed a large number of intermediate staining values. While CCND1 was indicated as a strong predictor of Uro vs GU subtypes, the large amount of overlap in staining intensities for this protein between these two subtypes may limit its utility for distinguishing them. Interestingly, the thresholds determined in parsimonious model building broadly correspond with the intersection points of curves in distribution plots (Fig. 5). This reflects on the robustness of the thresholds identified by the decision tree models, but also supports that these thresholds represent rational cut-off points for pathologic assessment.

GATA3 generally showed localized nuclear staining that was homogeneous across each core. However, proteins such as CCND1 and RB1 had greater heterogeneity in staining within a given core, and across paired cores for the same samples. Cytoplasmic KRT14 and KRT5 staining remained localized to the basal cell layer for low TCS values (TCS of 0-0.2) and was progressively more diffuse and homogeneous with increasing TCS values. For KRT14, the threshold range (0.64-0.68) indicates that the extension of expression beyond 2-3 cell layers from the basal epithelium is what determines the basal subtype. This contrasts slightly with KRT5, where the TCS threshold range falls slightly lower (0.09-0.34), indicating that almost any expression of KRT5 is enough to identify the basal subtype. Notably, p16 expression had a lower range of expression across all samples. The darkest staining intensity of p16 (representing a TCS of 1) was markedly less positive than expression for the other selected protein features.

## Discussion

In this work, we characterized patterns of protein expression and determined the accuracies of simple and complex decision tree models in classifying the molecular subtype of muscle invasive bladder cancers. Focusing on basal, Uro and GU subtypes which comprise more than 90% of MIBC cases identified GATA3, KRT5 and p16 as an attractive set of clinically available antibodies achieving 78% accuracy for the identification of Uro, GU and basal subtypes, and 91% accuracy for luminal/basal classification. Although limited data was available for testing, this work further identified a classification model using KRT14 and RB1 that could prove even more efficient and more accurate.

By exploring IHC expression patterns through clustering, correlation, and PCA analyses, we quantify the associations of these IHC features with each other and with IHC-based subtypes. These analyses highlighted the fundamental associations of the 24 proteins previously selected for subtype identification and their associations with subtype biology (1,14–16,21). As expected, these analyses showed the most similarity between Uro and GU clusters through their expression of luminal differentiation markers. As visualized in PCA analysis, Uro and basal subtypes showed greater overlap than GU and basal subtypes. This relationship is consistent with known features of these subtypes and interestingly, emphasizes how Uro and Basal features overlap in a subset of luminal tumors. This potentially reflects phenotypes such as that of the relatively aggressive Urobasal B subtype identified by mRNA profiling studies (7,14). Visualizations of IHC expression data by subtype clearly recapitulated aspects of subtype biology, indicating underlying genomic circuitry. The current work further quantifies these associations at the protein expression level, as indicated by positive correlative associations within luminal (GATA3, FOXA1, CDH3, PPARG) and basal (KRT5, KRT14) expression markers.

Consistent with previous work, we identify KRT5 and KRT14 as defining features of the basal subtype. Although we were unable to validate the accuracy of the model in a validation set, our parsimonious model building interestingly suggests the utility of KRT14 as a univariate classifier (Fig.3). This observation resonates with previous work identifying KRT14 as a defining feature of true basal cells in the bladder urothelium (29). Volkmer *et al.* identified various differentiation states in urothelial carcinoma which they characterize using IHC expression of KRT14, KRT5 and KRT20. They defined three differentiation states of bladder cancers, known as basal (KRT14+, KRT5+, KRT20-), intermediate (KRT14-, KRT5+, KRT20-) and differentiated (KRT14-, KRT5-, KRT20+) (29). In benign urothelial development, the same authors found that KRT14 expression marked the least differentiated, stem cell-like basal cells and occurred earlier than KRT5 and KRT20, which were found in intermediate cells and terminally differentiated umbrella cells, respectively. These changes in expression levels as a function of differentiation state may explain why both KRT5 and GATA3 are necessary to identify luminal and basal tumors, as KRT5 is identifying both intermediate and basal cell differentiation states, and therefore only classifies the basal subtype in the absence of GATA3. Comparatively, KRT14 alone may be sufficient to distinguish the basal subtype, as it distinguishes only the basal cell differentiation state.

Notably, much of the current literature highlights the utility of the same key differentiation markers described in these models and ROC analysis as candidate assays for subtype identification (10, 17–20). A number of studies propose a combination of two to four luminal and basal protein features such as FOXA1, GATA3, KRT5/6 and KRT14 (10, 19, 20) for bladder cancer subtyping. In this work, we show that the expression patterns of these protein features are highly overlapping, suggesting redundancy in using all of these assays for identifying subtypes. Further, binary ROC analyses demonstrate the comparable accuracies of these proteins for basal/luminal classification. The current work suggests that some of these previously published models may be simplified to a single protein feature, or one basal and one luminal protein feature.

GATA3 and KRT5 have specifically been proposed by multiple groups as the best set of features for luminal-basal classification (17, 18) with over 90% accuracy. Early work by Dadhania *et al.* (17) and a more recent study by Guo *et al.* (18) performed stepwise evaluations for determining the best set of IHC protein features for identifying basal and luminal subtypes (17, 18). Guo *et al.* distilled a large panel of genomic markers and key IHC proteins down to UPK2, KRT20, GATA3, KRT5 and KRT14 as defining IHC features of basal and luminal subtypes (18). They evaluated these key IHC features on whole slides and determined that GATA3, KRT14, and KRT5 showed the most consistent staining. Further logistic regression analyses determined GATA3 and KRT5 as the best pair of protein features. Further, the observations of both Guo *et al.* (18) and Dadhania *et al.* (17) demonstrated robust and reproducible staining patterns. Although, a notable difference with these studies is that they use IHC expression to determine mRNA subtype, not IHC-subtypes. Despite differences in subtype labelling and classifier building, we arrive at a similar conclusion with respect to the accuracy of a model using GATA3 and KRT5, complementing prior work and highlighting the robustness of these assays.

While a number of studies have focused on the identification of basal and luminal subtypes, to our knowledge, none of these efforts have been specific to the Lund classification. Particularly, none of these studies have expanded beyond basal/luminal classification to identify additional subsets of tumors within these subtypes. Increased granularity for identifying subsets of luminal tumors such as the GU subtype, which have distinct predictive prognostic associations, may be highly warranted. In MIBC, GU tumors have intermediate to poor prognosis (1, 8, 14). Additionally, the GU subtype, and tumors with similar features of genomic instability (luminal-unstable subtype in consensus model) have been associated with improved responses to neoadjuvant chemotherapy and immune checkpoint blockade therapies (11–13).

In the current study, we made classifiers to identify the GU subtype, which represents both a biologically and clinically distinct group. In models to identify basal, Uro and GU subtypes, the model using GATA3, KRT5 and p16 achieved 78% accuracy, comparable to that of KRT14 and RB1 at ~85% accuracy. While these findings with respect to models incorporating KRT14 and RB1 are only preliminary, they indicate that perhaps these two proteins may provide a simpler and more accurate classification. The multiclass model also suggested the use of RB1 to distinguish GU from Uro tumors, in agreement with both ranked proteins in ROC analysis and simplified model building (Fig. 4). This points to the utility of RB1 for identifying GU tumors, as well as CCND1 and TP63 as potential validation stains.

**Figure 4.**
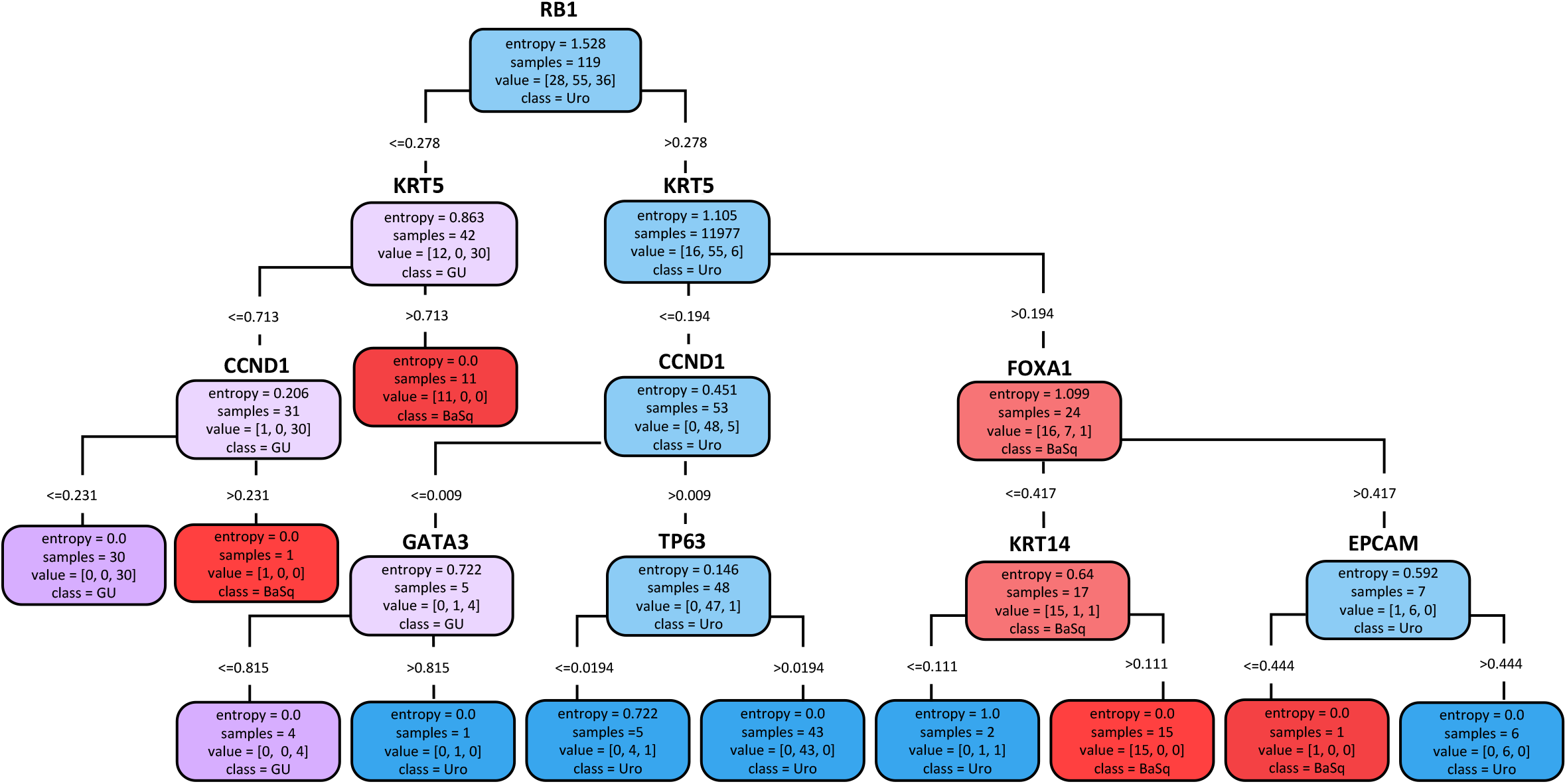
A multiclass decision tree classifier for MIBC subtypes achieves similar accuracy to simplified decision tree models. The model found to have optimal performance used RB1, CK5, CCND1, FOXA1, GATA3, TP63, KRT14 and EPCAM. Almost all final entropy scores were zero, indicating correct classification. As expected, the model had the greatest difficulty assigning Uro and GU subtypes in terminal leaves of the tree where entropy > 0.

**Figure 5.**
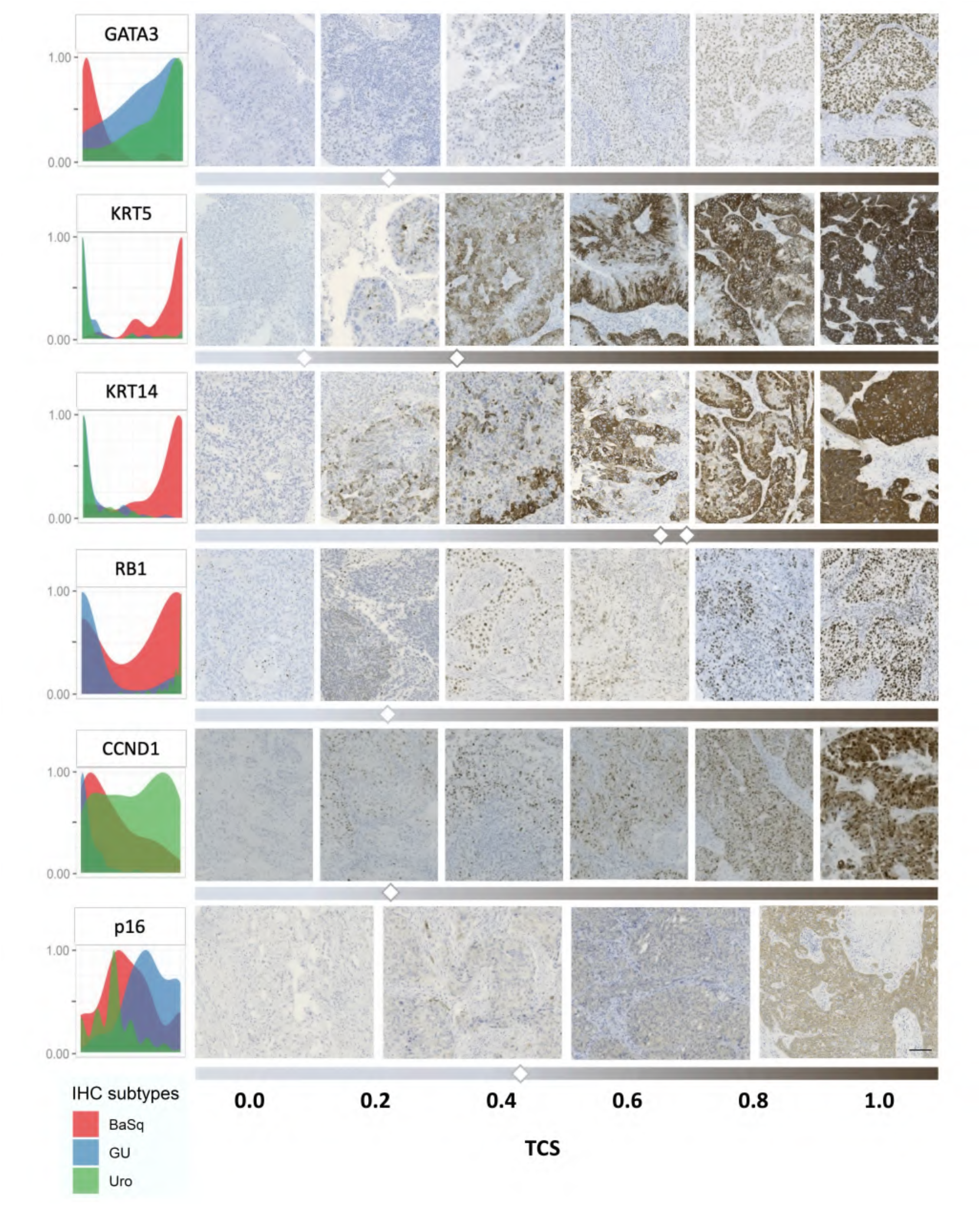
Subtype-specific expression patterns and distributions of key proteins highlight potential utility in classifying bladder cancers. Density plots (Left) show expression distributions for key proteins across samples of the basal (red), GU (blue) and Uro (green) subtypes. (Right)Visualization of IHC staining intensities and their respective TCS values. Representative staining for samples below and above threshold values are indicated by thresholding bars below IHC images.

Although KRT14 and RB1 may serve as useful protein features in subtype identification, an important consideration for these assays is their availability and reproducibility across clinical centres. Protein features which have not been validated across centres may show more variation in staining and result in issues of reproducibility. Thus, it is unclear whether the accessibility, and issues with reproducibility across centres may pose as a barrier to implementation of a model incorporating an assay for KRT14 or RB1. The initial model we proposed using GATA3, KRT5 and p16 was selected on the basis that these were key IHC features with assays that were also widely available in clinical laboratories. The same does not appear to be true for KRT14.

Aside from the clinical availability of these antibodies, another trade-off emerges between accuracy and simplicity when considering a more complex model optimized for accuracy. The multiclass model using expression of eight proteins (Fig. 4) achieved 82-86% accuracy, slightly improved from the GATA3 and KRT5 model (Fig. 2), and approximately equivalent to the model trained using KRT14 and RB1 (Fig. 3). While particular IHC features may be beneficial for clarifying subtypes, the multiclass decision tree classifier was much more complex, incorporating a larger set of features and nodes and therefore more opportunities for error. Although this model is optimized for accuracy, the trade-off is that this model becomes complex and uses far too many antibodies to be easily implemented, and that the addition of these extra proteins comes with limited benefit in terms of improving model accuracy.

Limitations of this work include the relatively small number of samples assessed once split into train and validation sets. Additional studies will be necessary to validate the accuracies of the models identified, and to investigate the utility of the identified cut-offs as visually assessable thresholds. We worked with previously assessed IHC expression data from tissue microarrays, but ultimately visual assessment of whole slides will be required to identify the extent to which intratumoral heterogeneity impacts these expression patterns. Further, although the clinically actionability of transcriptomic subtypes has been investigated, these must be explored specifically with respect to IHC-subtypes. Nevertheless, the simplicity of these IHC-based models compared to transcriptomic profiling provides opportunities for accelerating subtyping in both research and clinical practice. While this study was restricted to the MIBC setting, the provided datasets included non-muscle-invasive bladder cancers (NMIBCs) as well but provided too few samples to build separate models. Future work should assess the similarity of constructed models for subtyping classification in the NMIBC setting, as well as determine the utility of the models presented in this study.

## Conclusions

Altogether, this work clarifies the utility of GATA3, KRT5 and p16 IHC for identifying Uro, GU and basal subtypes, which represents the overwhelming majority of MIBCs. We demonstrate that a simple immunohistochemical decision tree classifier for determining these subtypes achieves ~78% to identify Basal/Uro/GU subtypes and >90% accuracy for distinguishing basal/luminal subtypes. While ultimately, the depth of a transcriptomic classification method may provide more comprehensive information than a surrogate IHC based classification, transcriptomics has not become part of routine clinical workup for bladder cancer. Here we demonstrate that simplified IHC assays can identify these subtypes with good accuracy. If clinical associations of these IHC subtypes are validated, they may hold important implications for prognostic or predictive stratification in the clinical setting.

## Acknowledgements

We thank the entire Lund bladder cancer group for their support of this work and use of their subtyping data from the 2017 and 2012 cohorts.

## Competing Interests

The authors declare no competing interests.

## Funding Statement

This study was conducted with the support of the Ontario Institute for Cancer Research through funding provided by the Government of Ontario (DMB). Additional support provided by the Cancer Research Society and Bladder Cancer Canada through the Operating Grant Funding Program (DMB).

## Author Contributions

CLJ and DMB conceived the manuscript which was written in conjunction with CH. CLJ and HG contributed to data analysis and figure generation. AS contributed to IHC evaluation and figure generation. GS conducted original data collection and to the conception of IHC-based profiling methods. RG contributed to the bioinformatic analyses. All authors contributed to manuscript editing.

## Notes

### Competing Interest Statement

The authors have declared no competing interest.

