## Supplemental Text and Tables for "Immunohistochemical assays for bladder cancer molecular subtyping: Optimizing parsimony and performance using Lund taxonomy"

### Supplemental Material

#### Supplementary materials and methods

IHC expression was reported using an intensity scale (0-3), and a percentage scale (percentage positive tumor cells, 0-9, in bins of 10%), or both (See IHC Feature Table). For CCNB1, RB1 and TP63, the intensity was disregarded and only the percentage of positive tumor cells was reported (range 0-9). For all remaining features, the intensity alone (range 0-3), or the intensity multiplied by the percentage of positive cells (range 0-3 x 0-0.9 = 0-2.7) was reported.

**Table S1**. Original scoring of IHC features and scaling across datasets.

| IHC Feature | *2017* | | *2012* | |
|---|---|---|---|---|
| | **Original assessment** | **Scaling** | **Original assessment** | **Scaling** |
| CCNB1 | Perc. | /9 | - | - |
| CCND1 | Perc.* Intensity | /2.7 | - | - |
| CDH1 | Perc.* Intensity | /2.7 | Intensity | Values could not be scaled to equivalent |
| CDH3 | Perc.* Intensity | /2.7 | - | - |
| CDKN2A(p16) | Intensity | /3 | - | - |
| CGHA | Intensity | /3 | - | - |
| E2F3 | Perc.* Intensity | /2.7 | - | - |
| EPCAM | Perc.* Intensity | /2.7 | - | - |
| FGFR3 | Intensity | /3 | - | - |
| FOXA1 | Intensity | /3 | - | - |
| GATA3 | Perc.* Intensity | /2.7 | - | - |
| KRT14 | Perc.* Intensity | /2.7 | Intensity | Values could not be scaled to equivalent |
| KRT20 | Intensity | /3 | - | - |
| KRT5 | Perc.* Intensity + staining pattern according to basal cell layer localization | Perc.* pattern/2.7 | - | - |
| NCAM1 | Perc.* Intensity | /2.7 | - | - |
| PPARG | Perc.* Intensity | /2.7 | Intensity | Values could not be scaled to equivalent |
| RB1 | Perc. | /9 | /9 | - |
| RXRA | Perc.* Intensity | /2.7 | Intensity | Values could not be scaled to equivalent |
| SYP | Perc.* Intensity | /2.7 | - | - |
| TP63 | Perc. | /9 | - | - |
| TUBB2B | Intensity | /3 | - | - |
| UPK3 | Intensity | /3 | - | - |
| VIM | Perc.* Intensity | /2.7 | - | - |
| ZEB2 | Intensity | /3 | - | - |

Perc. = Percentage; * = multiplication of indicated values; - = staining assessment did not differ between the two studies.

#### Assignment of tumor-cell phenotypes based on IHC data

Cellular phenotypes of advanced bladder cancer was defined by calculating *definition scores* as outlined by Lund (1,14):

| ***Subtype*** | ***Definition score calculation*** |
|---|---|
| **Uro/UroB:** | CCND1+, FGFR3+, RB1+, p16- |
| **Genom. Unstable:** | CCND1-, FGFR3-, RB1-, p16+ |
| **Mes-like:** | VIM+, ZEB2+, EPCAM-, E-Cad- |
| **Basal/SCC-like:** | KRT5+, KRT14+, GATA3-, FOXA1- |
| **Sc/NE-like:** | TUBB2B+, EPCAM+, E-Cad-, GATA3- |

Scores for each protein were normalized to the range-maximum, as indicated in Table S1. The definition scores were calculated by the Lund group (1,14), to assign tumor-cell phenotypes. Briefly, the highest positive score (>0.6) for any of the **Mes-like**, **Basal/SCC-like**, and **Sc/NE-like** subtype definitions classified these cases. For scores <0.6, the case was classified as **Uro/UroB** if the definition score was >0.6. If the case definition score is <0.6 it is defined as **Genomically Unstable** (this phenotype is thus defined as opposite to that of the **Uro/UroB**).

#### Supplementary data

**Table S3.** Ranked performance of individual proteins for identifying subtypes assessed using binary ROC analysis.

| | Uro vs. Basal | | GU vs. Basal | | Uro vs. GU | |
|---|---|---|---|---|---|---|
| Rank | **feature** | **AUC [95% CI]** | **feature** | **AUC [95% CI]** | **feature** | **AUC [95% CI]** |
| 1 | CK5 | 94.39 [91.25-97.52] | CK5 | 95.87 [91.81-99.93] | RB1 | 92.62 [87.17-98.06] |
| 2 | KRT14 | 94.11 [88.87-99.35] | KRT14 | 93.93 [88.63-99.22] | CCND1 | 90.11 [85.15-95.07] |
| 3 | CDH3 | 93.01 [89.05-96.98] | CDH3 | 93.74 [89.02-98.47] | p16 | 87.15 [80.93-93.37] |
| 4 | GATA3 | 91.46 [86.92-96.01] | FOXA1 | 89.48 [83.26-95.69] | TP63 | 81.68 [73.93-89.44] |
| 5 | FOXA1 | 91.32 [87.25-95.39] | GATA3 | 89.11 [82.05-96.17] | FGFR3 | 78.11 [70.4-85.82] |
| 6 | PPARG | 90.69 [87.03-94.35] | EPCAM | 88.98 [82.61-95.35] | CCNB1 | 71.86 [63.33-80.39] |
| 7 | EPCAM | 85.49 [79.75-91.23] | TP63 | 88.3 [81.27-95.34] | PPARG | 64.16 [54.19-74.13] |
| 8 | CDH1 | 77.26 [69.27-85.25] | PPARG | 86.45 [79.99-92.91] | RXRA | 63.45 [53.94-72.96] |
| 9 | KRT20 | 77.2 [72.47-81.93] | KRT20 | 79.38 [72.41-86.35] | UPK3 | 62.45 [55.04-69.87] |
| 10 | VIM | 71.15 [61.72-80.58] | CCND1 | 77.17 [67.54-86.8] | E2F3 | 62.13 [52.42-71.84] |
| 11 | TP63 | 67.77 [59.8-75.75] | CDH1 | 72 [61.54-82.45] | GATA3 | 60.09 [50.38-69.81] |
| 12 | CCNB1 | 65.67 [56.25-75.1] | p16 | 71.68 [60.98-82.38] | EPCAM | 58.04 [49.01-67.07] |
| 13 | UPK3 | 64.58 [59.67-69.48] | RB1 | 71.59 [61.86-81.32] | VIM | 57.42 [48-66.83] |
| 14 | FGFR3 | 62.78 [53.72-71.83] | VIM | 67.28 [56.39-78.16] | CDH1 | 56.87 [47.84-65.89] |
| 15 | CHGA | 56.25 [53.46-59.04] | E2F3 | 59.12 [47.54-70.71] | FOXA1 | 54.65 [44.8-64.5] |
| 16 | RB1 | 56.05 [46.51-65.6] | RXRA | 58.54 [47.03-70.05] | ZEB2 | 54.32 [51.24-57.4] |
| 17 | SYP | 52.21 [50.47-53.94] | UPK3 | 57.92 [51.56-64.27] | CK5 | 52.69 [43.49-61.89] |
| 18 | E2F3 | 51.91 [41.72-62.1] | CHGA | 57.41 [52.63-62.19] | CDH3 | 52.1 [42.11-62.09] |
| 19 | RXRA | 49.98 [40.27-59.7] | NCAM1 | 51.78 [42.61-60.96] | SYP | 52.07 [48.87-55.27] |
| 20 | p16 | 48.65 [38.69-58.62] | SYP | 50.93 [49.11-52.74] | KRT14 | 51.3 [41.88-60.72] |
| 21 | CCND1 | 48.46 [39.17-57.74] | CCNB1 | 46.27 [34.41-58.14] | TUBB2 B | 50.17 [43.66-56.68] |
| 22 | NCAM1 | 47.58 [40.05-55.11] | FGFR3 | 46 [34.96-57.04] | KRT20 | 47.69 [38.19-57.18] |

**Table S4.** Summary table of Basal/luminal decision tree four-fold validation and model accuracies

| Tree Number | Number of Branches | IHC Features Used | Feature #1- GATA3- Threshold | Feature #2- CK5 - Threshold | Accuracy |
|---|---|---|---|---|---|
| Tree 1 | 3 | GATA3 + CK5 | 0.21 | 0.16 | 0.93 (0.81-0.99) |
| Tree 2 | 3 | CK5 + GATA3 | 0.35 | 0.51 | 0.86 (0.73-0.95) |
| Tree 3 | 3 | GATA3 + CK5 | 0.21 | 0.34 | 0.95 (0.85-0.99) |
| Tree 4 | 3 | GATA3 + CK5 | 0.21 | 0.093 | 0.93 (0.82-0.99) |

**Table S5.** Summary table of Basal/GU/Uro decision tree four-fold validation and model accuracies

| Tree Number | Number of Branches | IHC Features Used | Feature #1- GATA3- Threshold | Feature #2- CK5 - Threshold | Feature #3 P16- Threshold | Accuracy |
|---|---|---|---|---|---|---|
| Tree 1 | 3 | GATA3 + p16 | 0.21 | | 0.42 | 0.74 (0.59-0.87) |
| Tree 2 | 3 | CK5 + p16 | | 0.51 | 0.42 | 0.72 (0.56-0.85) |
| Tree 3 | 3 | GATA3 + p16 | 0.21 | | 0.42 | 0.69 (0.53-0.82) |
| Tree 4 | 5 | GATA3 + CK5 + p16 | 0.21 | 0.19 | 0.42 + 0.50 | 0.76 (0.61-0.87) |
