## Supplemental Figures for "Immunohistochemical assays for bladder cancer molecular subtyping: Optimizing parsimony and performance using Lund taxonomy"

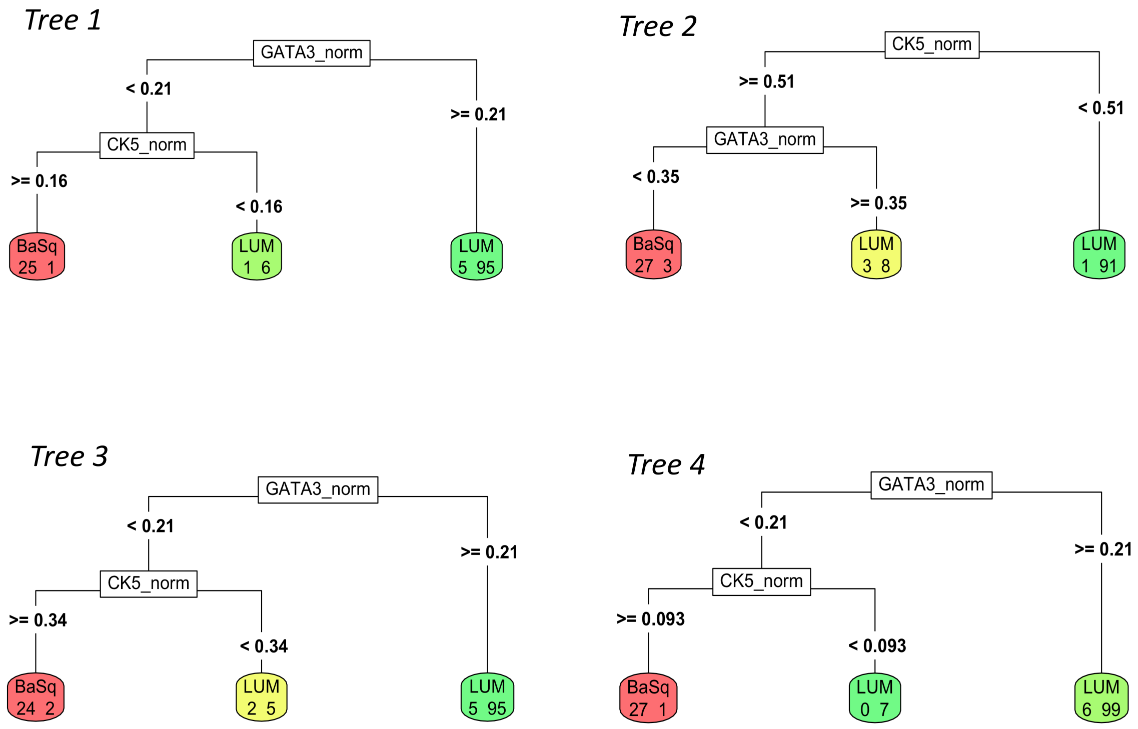


**Figure S1. Decision tree models in four-fold cross validation of Basal/Luminal model training using GATA3 and KRT5.**

### Seed 1

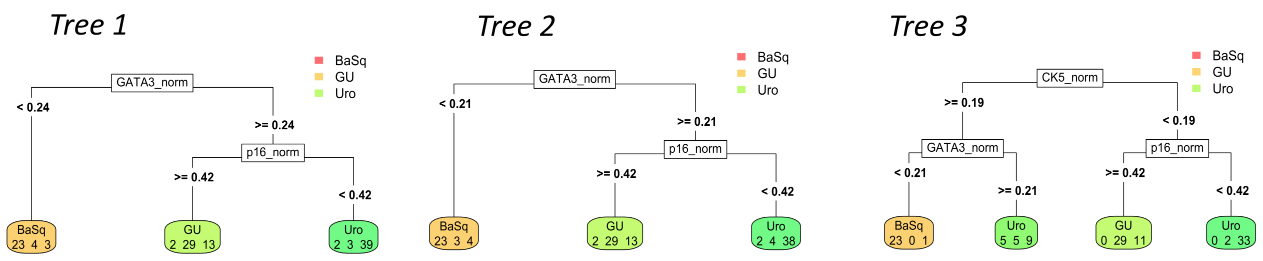


### Seed 2

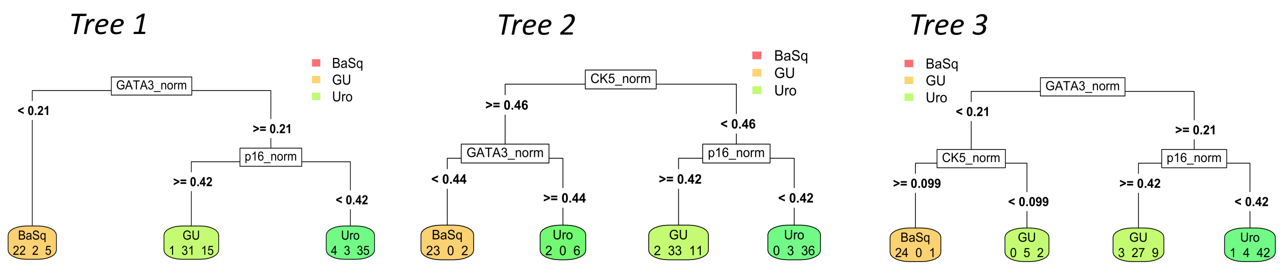


**Figure S2. Decision tree models in four-fold cross validation of Uro, GU and Basal subtype classification model training.**

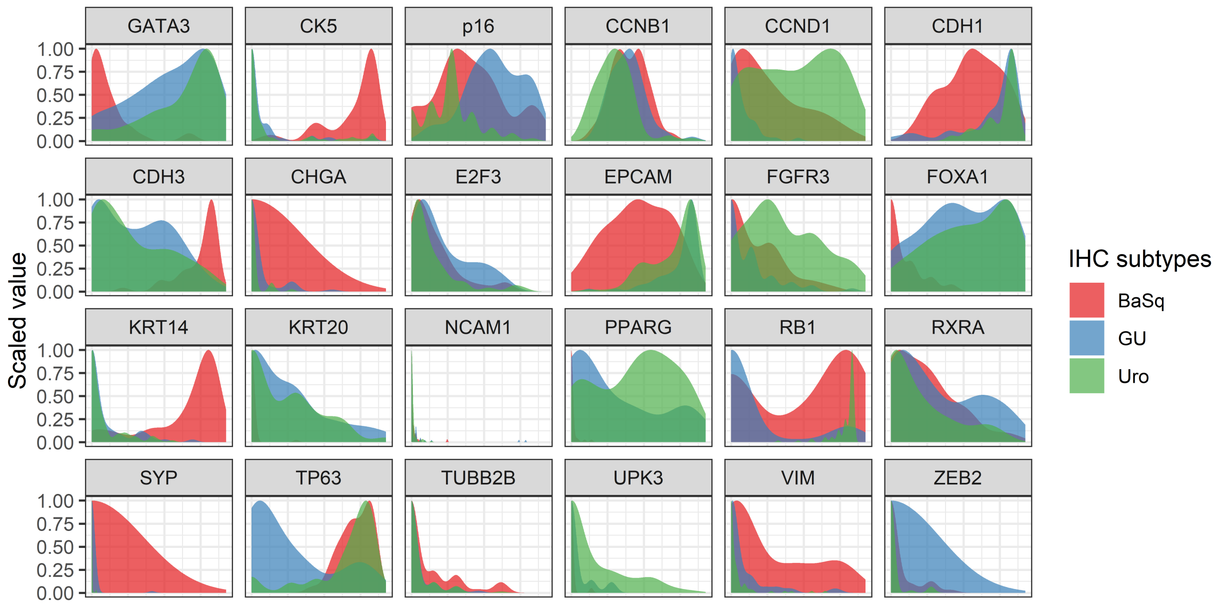


**Figure S3. Distribution of protein expression markers in Basal, GU and Uro subtypes.**
